# Coral hosts actively accumulate inorganic carbon for Symbiodiniaceae to maintain symbiotic relationships

**DOI:** 10.1101/2024.05.31.596804

**Authors:** Boya Zhang, Si Tang, Lu Liu, Meiting Xu, Yaqing Liu, Jianming Zhu, Weimin Xiao, Hongsheng Bi, Jin Zhou, Mark C. Benfield, Zhonghua Cai

## Abstract

High primary productivity of coral reefs is widely attributed to the mutualistic symbiosis between coral hosts and their microalgal partners (Symbiodiniaceae). Although the mechanisms maintaining this symbiosis have been considerable investigated, how the symbiont microalgae within coral get sufficient CO_2_ for photosynthesis still remains inadquately explored. Here, we hypothesized that corals may actively accumulate dissolved inorganic carbon (DIC) for microalgae to maintain the symbiosis. Carbon (HCO_3_^-^ and glucose) supply and consumption were evaluated in the scleractinian coral (*Goniopora lobata*) and its symbiont under light and dark conditions. Results suggest that Symbiodiniaceae were high DIC consumers, requiring about 2-3 fold more DIC than free-living species. The corals were high DIC producers, with internal concentrations up to 4.2 fold higher than in the surrounding seawater. In the absence of microalgae utilization, the excess DIC they produced appeared detrimental to their own growth. Moreover, transcriptomic analysis identified several DIC enrichment pathways are evolved in corals to attact the Symbiodiniaceae, including CO_2_ concentrating mechanisms, respiration, calcification and phosphoenolpyruvate carboxylase. Increased the CO_2_ dosage in seawater may induce coral symbiosis bleaching. Our findings can deeper reveal on the mechanisms sustaining coral symbiosis, and may help to predict how some corals respond to DIC imbalance under climate changes.

## Introduction

Coral reefs are recognized as having the highest levels of primary productivity (1500-5000 gC m^-^2a^-1^) in nature^1^, while growing in shallow oligotrophic waters^2,3^. The high productivity of coral reefs is widely attributed to their symbiotic relationship with the microalgal zooxanthellae^4,5,6^, recently reclassified as Symbiodiniaceae^7^. Within this mutualistic relationship, there exist complicated and intimate resource exchanges between these two partners. For instance, the microalgal symbionts can derive nutrients like inorganic carbon (C), nitrogen (N) and phosphorus (P) from corals^8,9,10^. In return, the corals can also acquire food such as glucose, amino acids, lipids, etc. by absorbing microalgal photosynthesis products even digesting parts of the symbiont population^11,12,13,14^.

From a stoichiometric perspective, coral’s high productivity requires a high organic carbon supply, this means that an active photosynthesis of the microalgal symbiont should be working. Functioning photosynthesis of Symbiodiniaceae requires relatively high level of dissolved inorganic carbon (DIC, CO_2_, and HCO_3_^-^ and CO_3_^2-^) supply, as Symbiodiniaceae posseses type II ribulose-1,5-bisphosphate carboxylase/oxygenase (Rubisco, the key enzyme for carbon fixation), which has lower CO_2_ affinity and specificity than type I Rubisco from other photoautotrophs^15,16^. Moreover, similar to other free-living microalgae that are usually challenged by insufficient DIC as CO_2_ diffuses 10,000 times more slowly in water than in air^17^, Symbiodiniaceae also faced this problem, which may be even worse as they inhabitated within the coral tissue and thus are physically separated from DIC in the surrounding seawater. It is therefore very interesting to undertand how Symbiodiniaceae can meet their CO_2_ requirement for photosynthesis under such a situation. Due to the close and symbiotic relationship between Symbiodiniaceae and the coral host, by now many efforts have been invested to study DIC interactions between these two partners^18,19,20^.

Understanding the DIC metabolism of the coral system is quite challenging because DIC within coral tissue is involved in several processes: (1) CO2 consumption by microalgal photosynthesis; (2) CO2 generation by the respiration of coral host and Symbiodiniaceae; and (3) CO2 generation and HCO_3_ ^-^ consumption due to coral calcification. Generally, besides HCO_3_ ^-^ in seawater, coral respiration is deemed to be the major contributor of photosynthetic CO_2_^21,22^, but different views have pointed out these CO_2_ will be rapidly transformed into HCO ^-^ and used by the host itself for calcification dependent on carbonic anhydrases (CAs) activities^23,24^, and it is thus believed that photosynthesis by microalgae and calcification by coral host compete HCO_3_^-^ in the tissue^25,26^. In contrast, others argued that these two processes are not in a competitive relationship, instead host calcification can supply CO_2_ for microalgal photosynthesis, as well as recognizing that photosynthesis can drive rates of calcification in the light^27,28,29^. More recently, researchers found that DIC concentrations inside corals exceed ambient levels and confirmed the presence of coral’s CO_2_ concentrating mechanisms (CCMs), nonetheless, the controversy between symbiont photosynthesis and host calcification on internal-DIC competition remains^24,30,31,32^. In addition, the productivity in Symbiodiniaceae may be carbon-limited even multiple DIC supply pathways exist in stable symbiotic systems^21,33,34^. The heterotrophic feeding of coral hosts also have a fairly important impact on symbionts^35,36^. There is much more to be studied and understood about the mechanisms supporting the complex coral symbiosis relationship.

Here, we explore the coral-microalgae symbiotic relationship focusing on the DIC supplies in the symbiotic system and propose that coral hosts may actively accumulate DIC for Symbiodiniaceae via respiration, calcification, CCMs, etc. To test this hypothesis, the scleractinian coral (*Goniopora lobata*) was employed as a model, which was widely distributed across the Indian and Pacific Oceans^37^. A series of carbon source (HCO_3_^-^ or glucose) utilization experiments were conducted under light or dark regimes to determine the carbon consumption characteristics of the coral host and symbionts, respectively. Transcriptomic analysis was then carried out to reveal corresponding molecular traits. Finally, CO_2_ enrichment experiments were performed to simulate the impact of high external DIC concentrations on coral symbiosis.

## Results

### DIC requirement in Symbiodiniaceae

To determine the photosynthetic DIC requirements of the symbiotic microalgae, the growth rates and carbon consumption rates of a monocultured Symbiodiniaceae (*Symbiodinium* sp.), isolated from the *G. lobata*, were evaluated at 2 mM, 4 mM, and 8 mM dosages of NaHCO_3_ over a period of 36 hours (Because N, P, etc. were still available in this period, the microalgae were not limited by other nutrients). In general, the symbiont showed the highest growth rate at 8 mM NaHCO_3_, which reached 0.53 d^-1^ and was 2.08 and 1.34 times higher than that of cells grown with 2 mM and 4 mM NaHCO_3_, respectively (Fig. 1a, Two-way ANOVA, F_2,6_ = 3.427, *p* < 0.05). As expected, faster-growing cells consumed more DIC. The average DIC consumption rates were 0.10 mM d^-1^, 0.18 mM d^-1^, and 0.26 mM d^-1^ corresponding to the 2 mM, 4 mM, and 8 mM NaHCO_3_ treatments, respectively (Fig. 1b), whereas, the DIC concentration from the blank control (without microalgae) remained almost unchanged (Fig. S1), suggesting the decrease in DIC was caused by microalgal photosynthesis but not by DIC run off. It was noteworthy that Symbiodiniaceae grown with different NaHCO_3_ concentrations all showed a linear increase in biomass with DIC consumption, suggesting that external DIC concentration is one of the limiting factors for its growth. Meanwhile, based on the slope of linear regression (0.012 mol g^-1^ for all treatments, Fig. 1b), it can be calculated that Symbiodiniaceae consume approximately 0.528 g of CO_2_ per gram of biomass growth, significantly higher than other free-living microalgae^38,39,40^. These findings suggest that available DIC is an important limiting factor for Symbiodiniaceae growth. Evidently, natural seawater with a DIC concentration around 2 mM is a carbon-deficient environment for Symbiodiniaceae^41^, so whether the DIC concentrations inside coral tissues can meet microalgal requirements and how this is achieved merits further investigation.

**Fig. 1:**
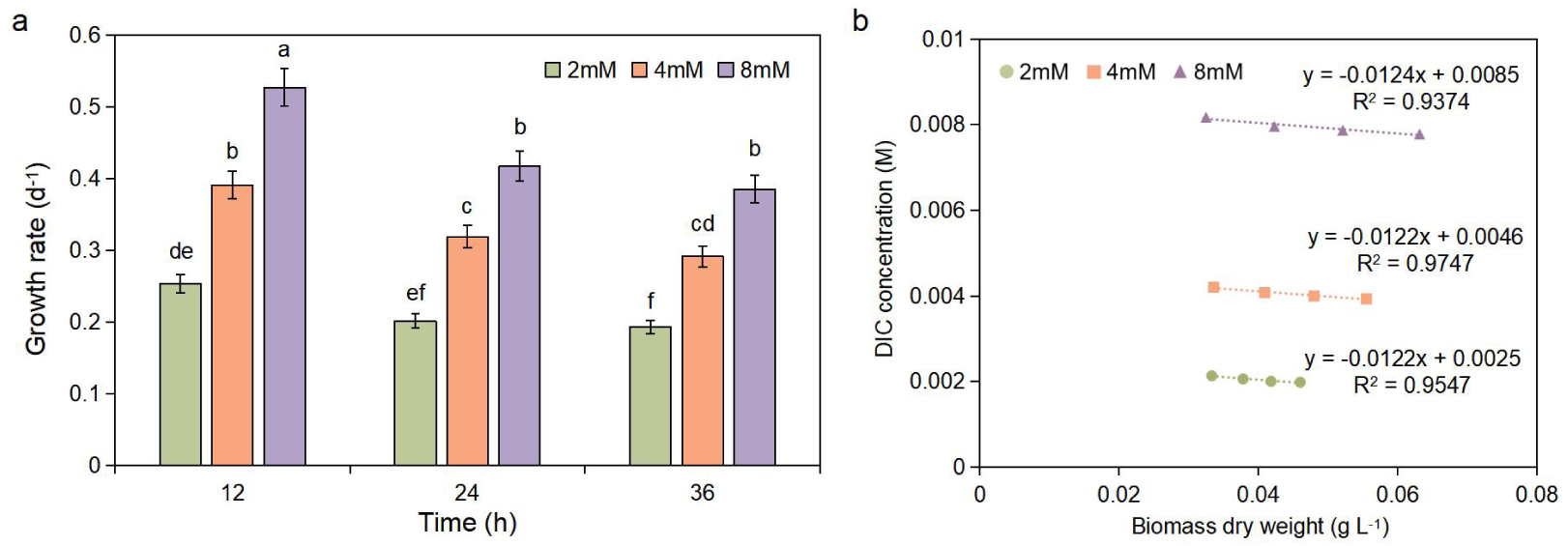
Growth rate and DIC consumption quantification of monocultured Symbiodiniaceae at different carbon concentrations. **a**, Growth rate of Symbiodiniaceae over a period of 36 h. The graphs show the mean ± SD (n = 3). Various letters above each bars indicate statistical significance (*p* < 0.05), calculated by two-way ANOVA with Tukey’s HSD posthoc analyses. **b**, Relationship between carbon consumption and biomass growth of Symbiodiniaceae (n = 3). Slopes indicate the carbon consumption for per gram of biomass growth. Data points are fitted with exponential decay functions. R^2^ values indicating the goodness of fits are shown.

### DIC production in the coral hosts

To explore the carbon budget within the coral-Symbiodiniaceae system, carbon supply experiments were performed under different light/dark regimes. Specifically, we had four treatments, i.e., DG (Continuous darkness and glucose as the only carbon source), DS (Continuous darkness and NaHCO_3_ as the only carbon source), LG (12 h/12 h light/dark and glucose as the only carbon source), and LS (12 h/12 h light/dark and NaHCO_3_ as the only carbon source). Here, CO_2_ or HCO_3_^-^ in the culture medium was pre-purged by the method of Yellowlees et al.^42^

Over the 8-day experiment, visual changes (images), DIC and glucose utilization of the corals were examined, respectively. Firstly, in the two treatments with glucose addition (LG and DG), we found that the length of coral tentacle extension differed in the light and dark, which were at most 1 cm longer in LG than in DG on day 4 (Fig. 2a). With regard to glucose comsuption, interestingly, the glucose was consumed by coral continuously in the light/dark regime, whereas corals grown in continual darkness stopped glucose uptake since day 4, suggesting a possible critical role of microalgal photosynthesis, which only occurs in the presence of light, in this process. At the end of the experiment (day 8), corals grown in LG and DG consumed 0.0723 g L^-1^ and 0.0404 g L^-1^ glucose, respectively (Fig. 2c, *p*<0.05). Notably, from day 4 onwards the internal-tissue glucose concentration of DG remained stable (approximately 0.0575 g L^-1^, Fig. 2e), but the DIC concentration continuously increased up to approximately 0.6009 g L^-1^(Fig. 2c). In contrast, the internal-tissue glucose concentration of LG kept increasing (from 0.0907 g L^-1^ to 0.1023 g L^-1^, Fig. 2e), and the DIC concentration gradually decreased (from 0.5632 g L^-1^ to 0.4297 g L^-1^, Fig. 2c). These results provided a good explanation for the morphological differences between corals in the LG and DG treatments: that corals could continuously utilize glucose in the light due to the consumption of internal-DIC by the symbiotic algae, whereas in the darkness, corals could not take up glucose when their internal-DIC reached a certain threshold. Secondly, in the NaHCO_3_ groups (LS and DS), the difference in DIC uptake of corals was also maximized between two groups on day 8 (LS=0.1180 g L^-1^, DS=0.0213 g L^-1^, *p*<0.05, Fig. 2b), suggesting DS absorbed few DIC in seawater. This is consistent with coral visual changes, the length of coral tentacles extended up to 1.3 cm in LS on day 8, but hardly extended in DS during the experiment (Fig. 2a). Moreover, we detected significantly higher DIC concentrations inside coral tissues compared to external seawater in both groups, reaching a maximum of 4.2 and 1.6 times that of seawater on day 4 in LS and DS, respectively (Fig. 2d). It is thus evident that DIC pools exist within coral tissues no matter microalgal photosynthesis is functioning or not. Notably, the enriched DIC within coral tissues in two glucose groups (LG and DG) were also found above, and the higher DIC was not being released into seawater, because the DIC concentration in external seawater of these two groups remained stable (approximately 0.02 g L^-1^, Fig. 2b).

**Fig. 2:**
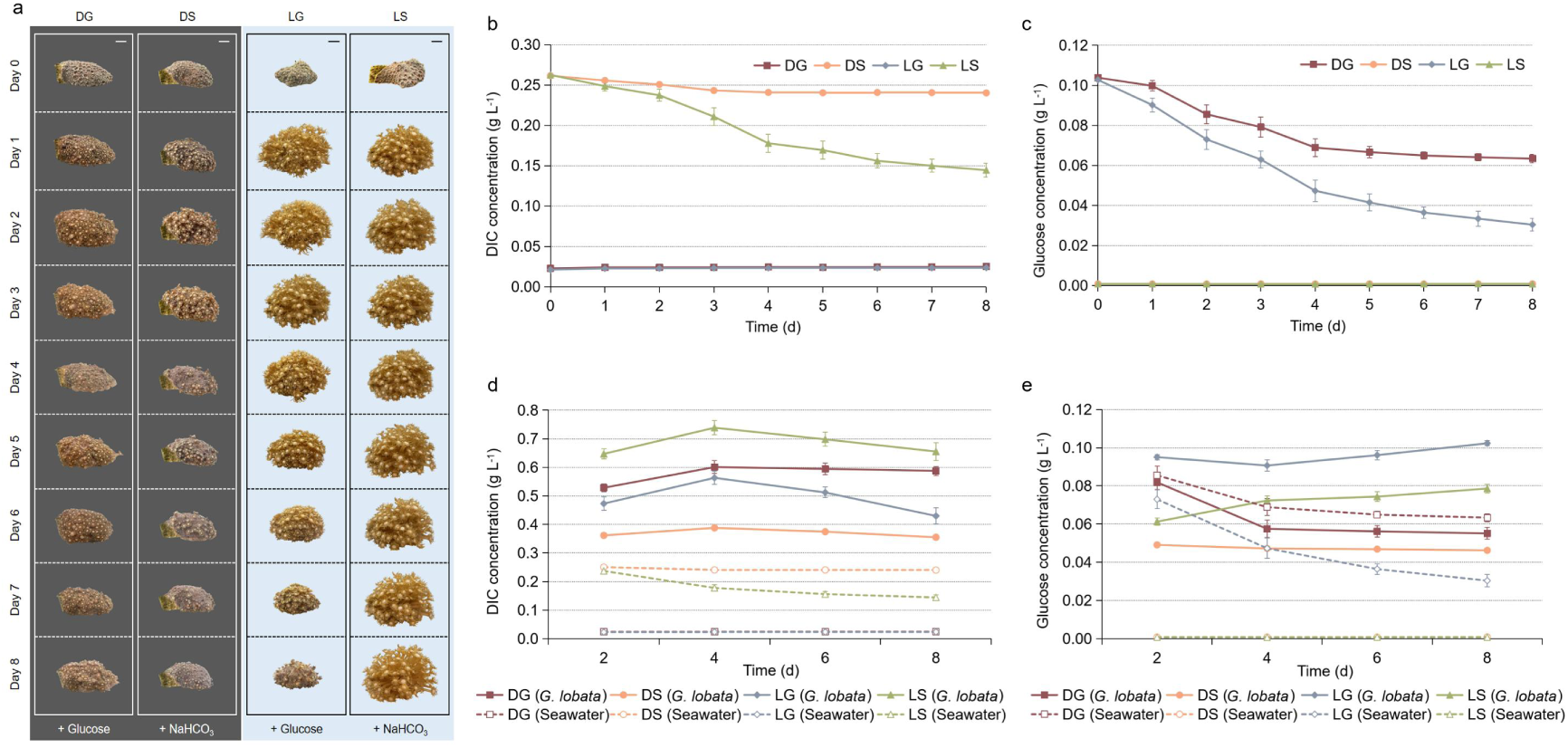
Variation of coral morphology, DIC and glucose concentration in seawater and corals under different carbon sources and light conditions. **a**, The visual changes of representative replicate samples of corals exposed to DG, DS, LG, LS. Scale bars, 1 cm. **b**, DIC concentration in seawater. **c**, Glucose concentration in seawater. **d**, DIC concentration in corals. **e**, Glucose concentration in corals. Straight lines and dotted lines indicate corals and seawater, respectively. The graphs show the mean ± SD (n = 3). DG indicates continuous darkness and glucose as the only carbon source. DS indicates continuous darkness and NaHCO_3_ as the only carbon source. LG indicates 12 h/12 h light/dark and glucose as the only carbon source. LS indicates 12 h/12 h light/dark and NaHCO_3_ as the only carbon source.

### Enzymatic activities and gene expression in carbon fixation

To resolve the origin of the high DIC pools in coral tissues, we analyzed the variance expressed genes (DEGs) of the coral host and Symbiodiniaceae, respectively. Coral grown in natural seawater (LS) was used as the control.

CAs, catalyze the interconversion of CO_2_ and HCO_3_^-^, and indicators of CCMs, were used to evaluate the DIC transformation of the coral symbiotic system. As to the enzymatic activity of CAs, LS (42.16 U g^-1^ prot^-1^) > LG (34.03 U g^-1^ prot^-1^) > DG (27.31 U g^-1^ prot^-1^) > DS (16.67 U g^-1^ prot^-1^) at day 8 (*p* < 0.05, Fig. 3a) CAs activity in two light groups (LS and LG) were gradually increased over culture period, but the two dark groups (DS and DG) showed a tendency to rise and then fall, showing that the CCMs of corals were affected by light. Subsequently, the expression of genes encoding different types of CAs was found in both coral hosts and Symbiodiniaceae. CAs are generally divided into α, β, and γ categories, with only α-CA in animals and all three categories in higher plants, microalgae, and cyanobacteria^43,44^. Notably, in coral hosts, seven α-CA genes were identified, and the DEGs of CA compared to LS were down-regulated in both dark groups (DG and DS), but up-regulated in the light group LG (Fig. 3d), indicating that their CCMs are associated with light. Moreover, in Symbiodiniaceae, different CA subtypes and locations may perform different functions. For example, the α-CA located in the cytoplasm and the β-CA1 in the chloroplasts were also up-regulated under light conditions (Fig. 3d), while the β-CA2 located in the cytoplasm was up-regulated in the dark groups (Fig. 3d).

**Fig. 3:**
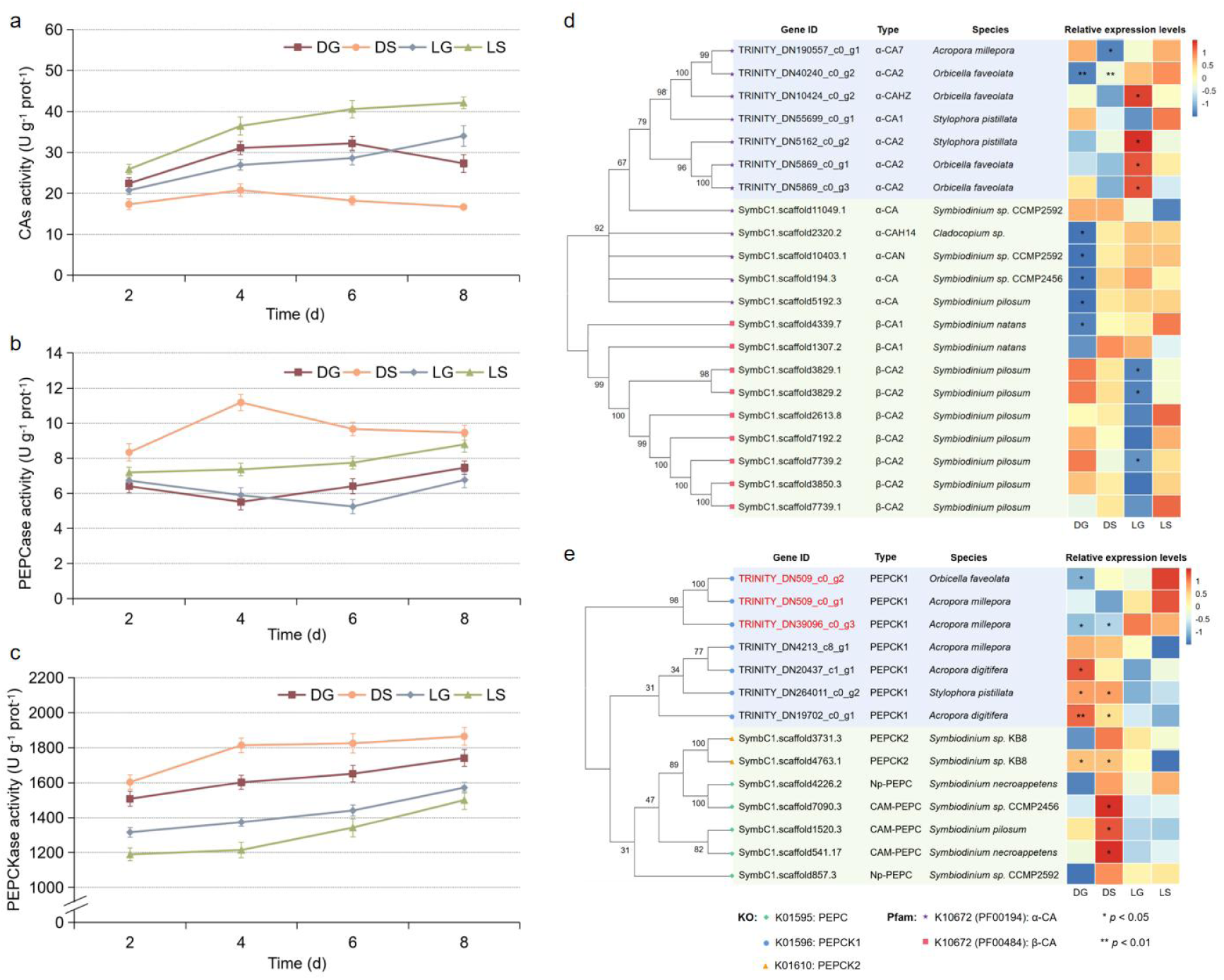
CAs, PEPCase and PEPCKase activities and gene expressions in coral hosts and Symbiodiniaceae. **a**, CAs activity in corals. **b**, PEPCase activity in corals. **c**, PEPCKase activity in corals. **d**, Differential expression of CA genes in coral hosts and Symbiodiniaceae. **e**, Differential expression of PEPC and PEPCK genes in coral hosts and Symbiodiniaceae. The graphs show the mean ± SD (n = 3). The gene sequences selected in the figure are more than 50% similar to the corresponding gene sequences included in the KEGG database. The phylogenetic tree was drawn by MEGA software, and the values indicate the branch lengths. Gene ID indicates the ID number of the sequence in the transcriptome on the comparison, beginning with TRINITY for host and SymbC1.scaffold for Symbiodiniaceae. Type indicates the specific type of the enzyme encoded by the genes, which was obtained through the KEGG module, Pfam, and Swissprot databases. Species indicate the similar species to which the sequence was compared in the NCBI-NR database. Relative expression levels are the log_2_fc values of each treatment group compared to the control LS. P indicates the significance of the difference between each treatment group and LS.

In addition, we also identified an interesting pattern on the phosphoenolpyruvate carboxylase (PEPCase) and phosphoenolpyruvate carboxykinase (PEPCKase), two key enzymes for the biochemical CCMs, i.e., C_4_ and crassulacean acid metabolism (CAM)^45^. PEPCase catalyses the conversion of phosphoenolpyruvate (PEP) to oxaloacetate and releases a CO_2_ around Rubisco in C_4_/CAM plants or supplements malate to the TCA cycle directly in all organisms^46^. PEPCKase catalyses the reverse pathway of PEPCase, converting oxaloacetate to PEP, and also plays a key role in gluconeogenesis^47^.

In terms of enzymatic activities, compared to LS, PEPCase was lower in the two glucose-added groups and higher in DS (Fig. 3b), while PEPCKase was more active in all treatments (especially in the two dark groups) (Fig. 3c). We attempted to explain the possible reasons by DEGs. Results showed that the PEPC genes were not found in coral hosts (Fig. 3e). Moreover, microalgal PEPC was mainly annotated as CAM types and highly expressed in DS. PEPCK genes in host and Symbiodiniaceae were generally up-regulated in the dark (Fig. 3e), which was consistent with the results of PEPCKase activities (Fig. 3c), indicating that the limitation of light would have forced both host and symbiont to synthesise glucose themselves. Notably, three sequences were up-regulated in coral hosts under light conditions, especially in LS (Fig. 3e-LS in red Gene ID), which was consistent with changes in PEPCKase activity at later stages (Fig. 3c-LS).

### Transcriptomic features of carbon metabolism of the coral host and Symbiodiniaceae

To explore the underlying mechanisms of DIC accumulation and transportation of the symbiotic system on a molecular level, we investigated the carbon-involved metabolisms in different coral treatments.

Glycolysis (purple arrows in Fig. 4c) and gluconeogenesis (red arrows in Fig. 4c) pathways were used as indicators for glucose utilization activities. In Symbiodiniaceae, compared to LS, all 9 genes annotated in glycolysis were down-regulated in DG, DS, and LG, and the genes encoding three rate-limiting enzymes of glycolysis, glucokinase (GLK) / hexokinase (HK), 6-phosphofructokinase (PFK), and pyruvate kinase (PK)^48,49^, were 2 out of 3 significantly down-regulated (log_2_fc > 1, *p* < 0.05) in DG. These lines of evidence pointed to external glucose being poorly utilized by Symbiodiniaceae, as was also evident from its DEGs of photosynthesis (Fig. S2). Meanwhile, the genes encoding the rate-limiting enzymes of gluconeogenesis, PEPCK2 and fructose-1,6-bisphosphatase (FBP)^50^, were up-regulated in all three treatments, even significantly up-regulated (log_2_fc > 1, *p* < 0.05) in DG, suggesting that it was consuming its own fatty acids to generate glucose.

**Fig. 4:**
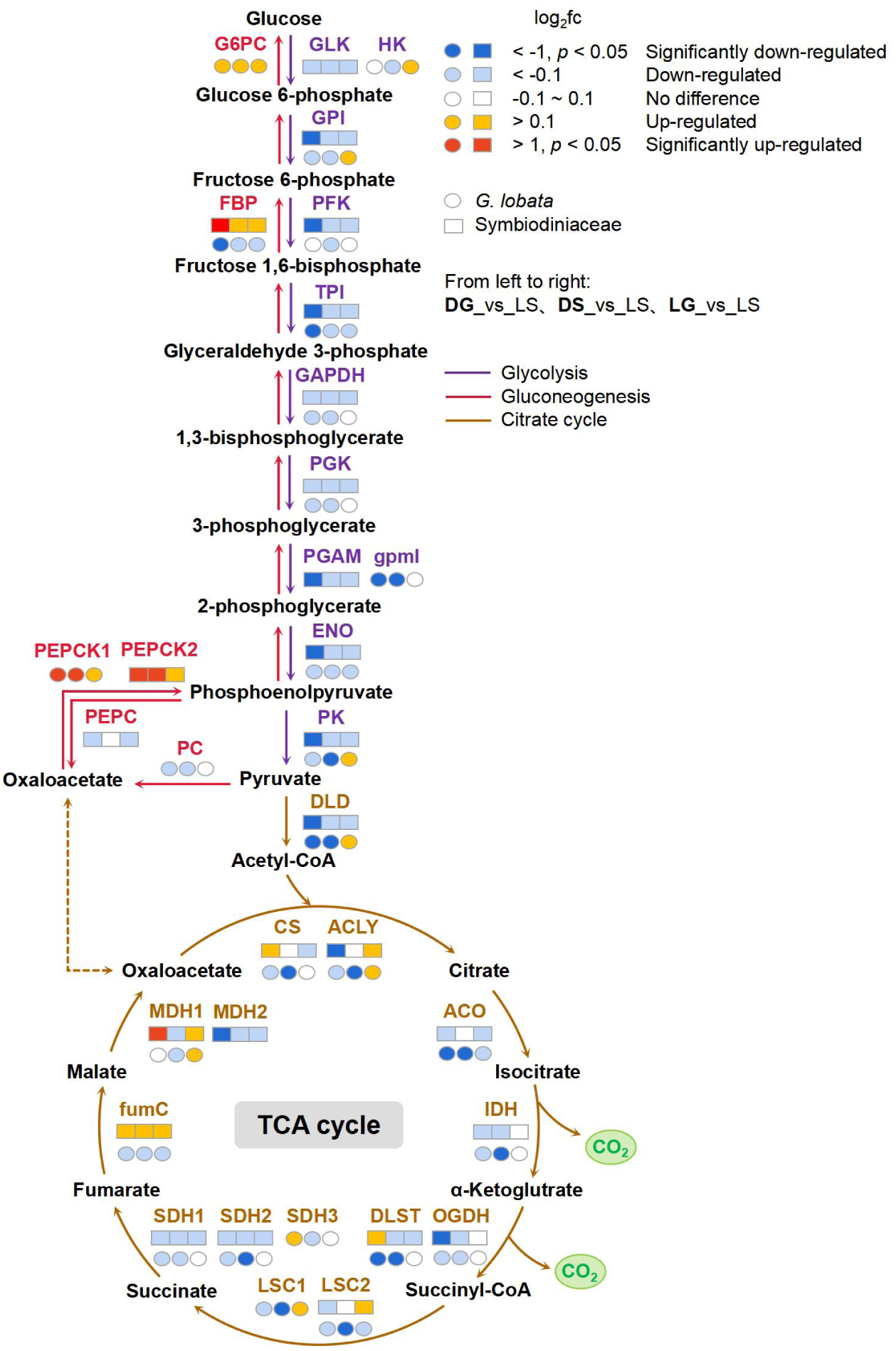
DEGs of glycolysis, gluconeogenesis and TCA cycle genes in coral hosts and Symbiodiniaceae. The expression results of one sequence with the largest length of the transcriptome compared with the reference sequence of the KEGG database were selected for each gene in the figure. ○ indicates coral host, and □ indicates Symbiodiniaceae, where the colors indicate the log_2_fc values of each treatment group compared to the control LS, from left to right once for DG, DS, and LG. Heatmap colors indicate relative expression levels, also the log_2_fc values of each treatment group compared with the control LS. *p* indicates the significance of the difference between each treatment group and LS.

In contrast, in coral hosts, out of 9 glycolysis-related DEGs, although 7 and 9 genes were down-regulated in DG and DS, respectively, but 7 genes (including three rate-limiting genes) were unchanged or up-regulated in the light group LG, which is in accordance with the result from Fig. 2c-LG. Unsurprisingly, partial gluconeogenesis-related genes (e.g., PEPCK1) of the host were up-regulated in DG, DS and LG, probably due to the external glucose decrease in the post-culture period. The TCA cycle (brown arrows in Fig. 4c), which is the central pathway for CO_2_ production in aerobic organisms, was further explored. In Symbiodiniaceae, compared to LS, the TCA cycle generally exhibited a decreased pattern, suggesting a deceleration in respiration. Specifically, 9, 8 and 7 out of 13 genes annotated in the TCA cycle were down-regulated in DG, DS, and LG, respectively. As to DEGs in coral hosts, almost all genes were down-regulated in both dark treatment groups. Conversely, 10 out of 14 TCA-related genes were unchanged or up-regulated in LG, indicating that the host accelerated respiration to produce CO_2_ in the light, combined with the carbon consumption in coral tissues from Fig. 2d-LG. This CO_2_ was likely to be supplied to the microalgae.

To further investigate whether CCMs from coral hosts acts on microalgal photosynthesis or on its own growth, calcification, the coral-specific DIC metabolic pathway central to coral physiology and health^51^, was also analyzed. In general, calcification-related genes displayed a decreased trend under dark conditions and an increased trend under light conditions. Specifically, compared to the no-glucose added group (LS), out of the 88 ion transporters DEGs responsible for active transportation, i.e., calmodulin (CaM), calcium-binding proteins (CaBPs), Na^+^/H^+^ transports, HCO_3_^-^ transports and CAs^52,53^, 71 and 85 displayed a down-regulation in DG and DS, respectively (Fig. 5a), for a total decrease of 59.31-log_2_fc and 77.57-log_2_fc (Fig. 5b). In contrast, the expression of DEGs in LG was 58 out of 88 genes unchanged or up-regulated (Fig. 5a), for a total increase of 20.05-log_2_-fold (Fig. 5b), suggesting a correlation between coral calcification and light, rather than glucose. Furthermore, galaxins, the sole skeletal organic matrix proteins definitively identified in corals^54^, continued to show a consistent down-regulation in DG and DS, while exhibiting a 2.19-log_2_-fold up-regulation in LG (Fig. 5b). The DEGs of other skeletal organic matrix proteins associated with early calcification^55,56,57,58^, were over 90% down-regulated as well in two dark treatments, but 50% unchanged or up-regulated in LG (Fig. 5a), which provides further evidence of the predominant influence of light conditions, rather than carbon sources on calcification.

**Fig. 5:**
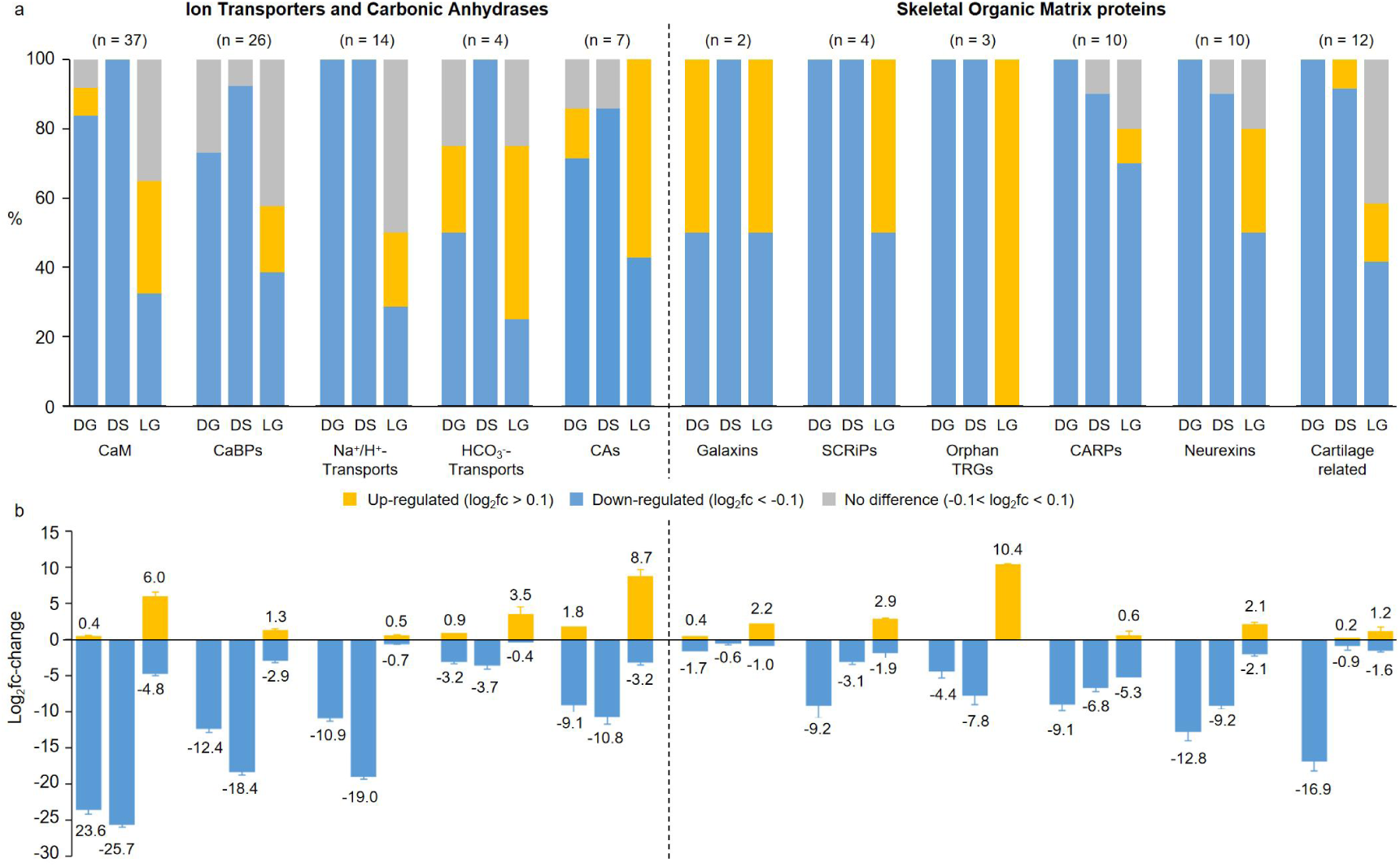
Comparison of differential expression of genes related to calcification in coral hosts. **a**, Summary of changes on calcification genes in different categories (ion transporters, carbonic anhydrases, and skeletal organic matrix proteins) for DG, DS and LG treatments. Column colors indicate log2fc values for each treatment group compared to control LS, with yellow up-regulated, blue down-regulated, and grey non-regulated. The total numbers of transcripts in each category are indicated above the corresponding bars. **b**, Summary of the corresponding mean log fold-change values for each of the classes of genes responding in part a. Actual fold-change values are indicated above or below each bar.

### Elevated external CO_2_ not low pH, induced detachment of Symbiodiniaceae

To examine the universality of our hypothesis that the corals may actively concentrate their internal DIC pool to favor symbiont photosynthesis as a primary evolutionary strategy for “retention” of symbiont microalgae, two scleractinian corals (*G. lobata* and CO_2_ mixture gas continuously aerated, pH = 7.0), HCl group (HCl added, pH = 7.0) and control group (natural seawater, pH = 8.2). Results showed that in CO_2_ group, some corals lost more than half of their symbiont population, resulting in a bleached appearance(Fig. 6ac). In the HCl group with the same pH, all corals maintained normal symbiont populations (Fig. 6a), suggesting that higher external CO_2_ levels instead of other factors play an decisive role in the loss of the corals’ symbiotic microalgae. Specifically, the chlorophyll of *G. lobata* and *A. digitifera* decreased to about 50% of that in the control groups on day 12 and day 3 in CO_2_ groups (Fig. 6b). We further explored whether the CO_2_ solubility impacted on the microalgae loss. In this study, the relationship between bubble diameter and CO_2_ solubility was determined by a Planktonscope and a total organic carbon analyzer (Fig. S3abc). Results showed that smaller bubbles (d_32_ = 467 in group 2, Fig. S3abc) in the microbubble plates led in higher CO_2_ solubility (0.1703 g L^-1^ and 0.2000 g L^-1^ in group 2, Fig. 6c), and resulted in higher microalgal loss rates (59% and 84% in group 2, Fig. 6c). To determine whether stress impacted the coral bleaching, anti-oxidative enzymes, i.e., superoxide dismutase (SODase) and catalase (CATase) activity were detected in coral holobiont. Compared to natural conditions, the CO_2_ group of two corals had similar activities of SODase and CATase (*p* > 0.05), but the HCl group significantly decreased the activities of the two enzymes (Fig. 6de, Two-way ANOVA, *p* < 0.05). Meanwhile, the photosynthetic activity (Fv/Fm, Fig. S3de) of detached Symbiodiniaceae in seawater was similar to that of the symbiotic Symbiodiniaceae and possessed higher CAs activity (57.37 U g^-1^ prot^-1^ and 46.32 U g^-1^ prot^-1^, Fig. 6f) in two corals, suggesting that the departure of Symbiodiniaceae from the host was not caused by stress response of excess CO_2_ and they survived well outside of the corals.

**Fig. 6:**
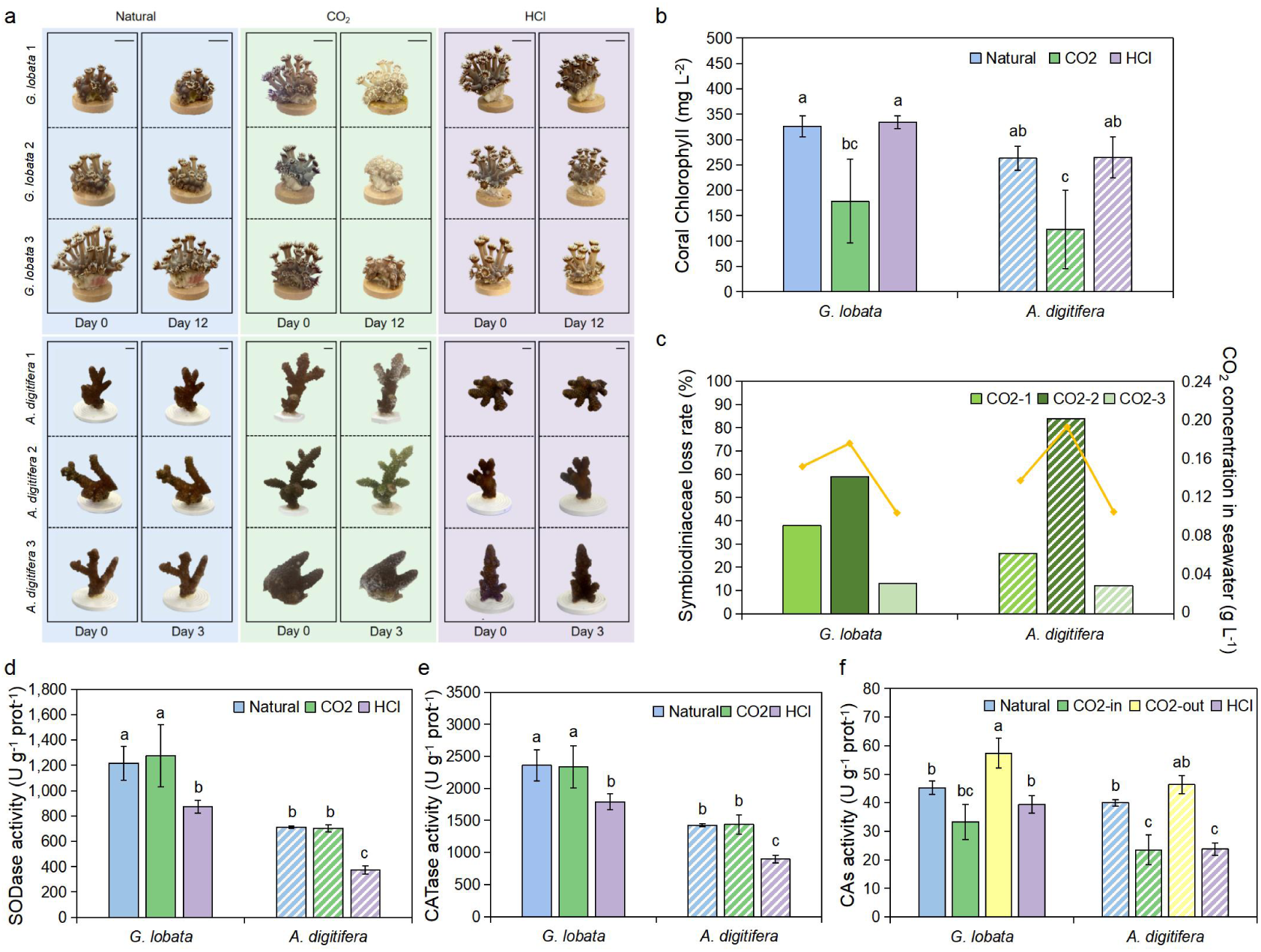
Characterization of *G. lobata* and *A. digitifera* after external CO_2_ and HCl addition. **a**, Visual changes of corals. **b**, Chlorophyll of corals on the last day. **c**, Symbiodiniaceae loss rates (bar charts) and CO_2_ concentration in seawater (line graphs) of three CO_2_ treatments on the last day. **d**, SODase activity of corals on the last day. **e**, CATase activity of corals on the last day. **f**, CAs activity of Symbiodiniaceae on the last day. Natural indicates natural seawater control group. CO_2_ indicates 1% CO_2_ mixture gas continuously aerated group. HCl indicates pH reduction with HCl addition group. The graphs show the mean ± SD (n = 3). Various letters above each bar indicate statistical significance (*p* < 0.05), calculated by two-way ANOVA with Tukey’s HSD posthoc analyses.

## Discussion

Coral reefs are highly productive ecosystems that thrive in oligotrophic waters. The success of this Darwinian paradox is mainly attributed to the successful symbiotic relationship between the coral host and the symbiont Symbiodiniaceae^59,60^. Evidently, efficient microalgal photosynthesis is paramount for this symbiotic system. However, Symbiodiniaceae possess type II Rubisco enzymes characterised by a low CO_2_ affinity, and they may also have higher CO_2_ demands^15,16^. Here, we specifically quantified the carbon requirements of isolated symbiotic microalgae, which consumed about 0.528 g CO_2_ per gram of biomass grown (Fig. 1b), suggesting that the Symbiodiniaceae are “high consumers” of DIC. This is a far higher DIC demand than that of some other free living microalgae. For example, *Haematococcus pluvialis* consumes 0.308 g CO_2_ for per gram of biomass growth^40^, *Spirulina platensis* requires 0.256 g CO_2_^38^, and *Chlorella vulgaris* requires requires 0.218 g CO_2_^39^. Obviously, the symbiont microalgae must somehow cope with DIC limitation and perform photosynthesis successfully and efficiently. It is reasonable that they may find high DIC sites or habitats within coral tissue.

In this study, we employed *G. lobata* as model, and found that these scleractinian corals are “high producers” of DIC, possessing a high internal-DIC pool up to 4.2 times higher than the DIC concentrations of external seawater (Fig. 2d-LS). This result supports carbon budget in internal symbiosis in Xu et al.^20^ We further discovered that the DIC pool remained when no external DIC was provided (Fig. 2d-DG/LG), showing that it does not depend on external seawater DIC and was more associated with its metabolic CO_2_ production. More interestingly, the DIC pool was rarely consumed in the dark (Fig. 2d-DG/DS), which means although the calcification consumes DIC, the coral hosts were unable to efficiently eliminate CO_2_ without symbiont photosynthesis. This result differs from the explanation that internal-DIC pool in corals is used for photosynthesis and calcification^30,31,32^, supporting the Furla et al.^24^ view that the host DIC is actively supplied to microalgal symbionts in a light-dependent manner and not used as a source for calcification. In addition, we found the significance of symbiont DIC removal to the host. The corals can continuously consume external glucose only in the light (Fig. 2c-LG), whereas they cannot sustainably utilize glucose in the dark (Fig. 2c-DG). Based on changes in the host tissues DIC (Fig. 2d), we presumed that if there is no photosynthetic consumption, the corals could not take up glucose when their internal-DIC reached a certain threshold. Pogoreutz et al.^61^ showed that the addition of sugar (including glucose) readily attributed to a fourfold increase of coral-associated microbial nitrogen fixation and then induced bleaching. In our experiment, the coral did not bleach during 8 days, but it is undeniable that further studies should consider the effects of N, P, etc. more comprehensively. Since we know that the symbiont microalgae require high levels of DIC for photosynthesis, the coral hosts, having a low DIC removal capacity, accumulate DIC in the light seemingly a detrimental behaviour for themselves, but actrually would be beneficial or even necessary to maintaining the host-Symbiodiniaceae symbiosis.

Results from the transcriptome support and explain our hypothesis that the DIC pool accumulated by corals may be the basis of symbiosis on a molecular level. Foremost, we found that the coral have evolved several DIC enrichment pathways, and they mainly serve the photosynthesis of the microalgae. Although several studies have explored CCMs in coral hosts^20,62,63^ and some of them proposed that the host’s CAs are involved in calcification^53,64^, our results provide evidence that the host CAs do not act primarily on their own calcification or emission of respiration CO_2_, because they have relatively low CAs activity in the dark compared to the light (Fig. 3a). The CAs are most likely functioning to supply DIC to the microalgal photosynthesis. This fully explains why coral hosts, having taken up enough DIC (Fig. 2d), continue to enrich their quantities of it. More detailed CA annotation results indicate that coral host-symbiont may possess different CA types with distinct functions and enhance carbon synergistically (Fig. 3d). Similarly, Tansik et al.^65^ also found that the supply of DIC in the coral symbionts is usually under the control of the hosts. If that is true, there may be other pathways that have evolved to “retain” the Symbiodiniaceae during coral symbiosis that have not been discovered yet.

More interestingly, we have identified a potential carbon fixation pathway within coral hosts that could supplement the CCMs, i.e., the PEPCK pathway. Notably, PEPCKase can act as a carboxylase in invertebrates^66,67^. In natural conditions, when the symbionts lacked DIC at the late stage, CAs activity gradually weakened while PEPCKase activity gradually increased. At the same time, the gene expression of three PEPCK sequences was significantly up-regulated (Fig. 3e-LS in red Gene ID), which is different from the role played by PEPCK in gluconeogenesis (Fig. 3e-LS in black Gene ID). In this process, PEPCKase may serve as PEPCase, converting PEP into oxaloacetate, which efficiently concentrates DIC in the C_4_ pathway, just as the microalgal C_4_ pathway has been reported^68,69,70^. There is another possibility that oxaloacetate formation by PEPCKase replenishes the TCA cycle under increased malate efflux, which accelerates tissue carbon flux to meet the photosynthetic requirements of the microalgae. This mechanism bears similarities to the increasing number of microalgal studies in recent years^71,72,73^. Notably, in the case of corals, the hosts play a pivotal role in facilitating this step of carbon fixation for the symbiont.

Furthermore, CO_2_ produced by host respiration is an important carbon source for Symbiodiniaceae photosynthesis. When glucose is used as the only carbon source, the hosts’ glycolysis genes (purple arrows in Fig. 4c-LG) and TCA cycle genes (brown arrows in Fig. 4c-LG) were up-regulated in the light, which showed that they were uptaking glucose for metabolic needs to release by-product CO_2_. These findings have similarities with previous studies that heterotrophic feeding and glucose enrichment can elevate the respiration rate of coral hosts, subsequently enhancing symbiont photosynthetic rates^74,75^. What’s more, the hosts must prevent more CO_2_ production without microalgal photosynthesis, i.e., by stopping the uptake of external glucose and accelerating gluconeogenesis to meet its own metabolic needs (Fig. 4c-DG). All the above corroborate our physiological indicators (Fig. 2cde-DG, LG).

Surprisingly, our results suggest that coral calcification is more likely to be a provider of DIC to microalgal photosynthesis rather than a competitor for DIC. Previous experiments shown that calcification is a competitor for photosynthesis, consuming 60% of the respiration-based CO_2_^25,26^. In contrast, it has been argued that there is no competition, and calcification rates are higher under light conditions compared to darkness^27,28,29^. Numerous hypotheses have been proposed to explain the mechanisms behind light-enhanced calcification. The most widely supported hypothesis suggests that the photosynthetic metabolites produced by microalgae can supply ample nutrients for host growth, thereby accelerating the calcification process^76,77,78^. However, in our experiment, even with sufficient external nutrition, all coral calcification genes were down-regulated when exposed to darkness (Fig. 5-DG), and up-regulated or unchanged in the light (Fig. 5-LG), suggesting that the correlation between calcification and coral growth is not robust. This is consistent with experimental results under dark conditions where both HCO_3_^-^ in external seawater and DIC in coral tissue remained utilized (Fig. 2b,d-DS), indicating that calcification may not be an actively-driven process. It can be inferred that coral calcification doesn’t occur in tandem with host growth but is primarily fueled by the strong carbon demands of symbiotic organisms. The CO_2_ released during calcification isn’t promptly expelled from the coral body but is instead absorbed and utilized by Symbiodiniaceae, which may provide a new explanation for the controversial issue of the coral as a carbon source or carbon sink^79,80^.

Finally, to confirm the significance of DIC in maintaining the coral-Symbiodiniaceae relationship, we performed 1% CO_2_ gas dosing addition experiments to increase the DIC concentration by a factor of 1.2 times higher than the natural seawater. Notably, compared to HCl addition group with the same pH, *G. lobata* and *A. digitifera* lost their Symbiodiniaceae on day 12 and day 3, respectively (Fig. 6ab), suggesting it was high CO_2_ rather than low pH that caused the microalgae to depart from the coral. Meanwhile, the microalgal loss rates were positively correlated with CO_2_ solubility (Fig. 6bc), suggesting that the higher external CO_2_ concentration was more attractive to the microalgae. The enzymes associated with anti-oxidadation activities of corals were not increased in CO_2_ group (Fig. 6de), and more interestingly the detached microalgae in seawater had higher CAs activity (Fig. 6h) and similar photosynthetic efficiency (Fig. S3d,e) than those within corals. This suggests that the coral bleaching in our experiments was not a result of the coral host expelling Symbiodiniaceae because they were not functioning properly, but most likely a result of the microalgae actively departing from the coral. From an evolutionary biology view, the microalgae’s behavior is paradoxical, implying that they actively abandon the protection of their hosts. However, our experiments dramatically increased the CO_2_ concentration in a short time, which could have avoided the limitation of other nutrient factors and made the microalgae respond more quickly. Another study found a similar phenomenon that the increased CO_2_ caused stress and reduced Symbiodiniaceae in *Galaxea fascicularis* and caused bleaching^81^. Undeniably, there are many resources that influence the coral-Symbiodiniaceae symbiosis^8,9,10,11^, but our experiment provides good evidence demonstrating the importance of actively DIC accumulation by the hosts in maintaining coral symbiotic relationships, because once the external DIC reaches concentrations sufficient for Symbiodiniaceae growth, the microalgal may leave the host.

In summary, in this study, we provide a new perspective to understand the basis of coral symbiosis. We propose that the coral hosts may have complementary DIC utilization characteristics with Symbiodiniaceae and actively accumulate DIC for the symbiont microalgae. The internal-DIC in coral tissue is supplied to Symbiodiniaceae, not used for calcification. Coral hosts CCMs, respiration and calcification are all critical for the accumulation of DIC for microalgal photosynthesis, and there are even other possible carbon fixation pathways, i.e., PEPCK (Fig. S4). Furthermore, our hypothesis argues that the coral bleaching caused by heatwaves and acidification^82,83,84^ may not be comprehensive, because it is also possible that as more greenhouse gases are incorporated into the ocean, it results in a situation where the hosts can no longer provide a relatively high CO_2_ environment, and thus lose the attraction to the microalgae, resulting in the collapse of the symbiosis. This may provide a new perspective on coral bleaching under climate change. Undoubtedly, to confirm these hypotheses, further experimental efforts over extended periods are needed, as well as consideration of the interactions with other available nutrients.

## Methods

### Coral collection

The colonies of scleractinian corals (*G. lobata*) were collected in Dapeng Bay (22°59’N, 114°33’E), Shenzhen, People’s Republic of China in September 2022. The coral colonies were collected at a water depth of ∼5 m and divided into 5×2×2 cm^3^ blocks using a Gryphon AquaSaw XL (American) and then were immediately transported to the indoor aquaria facilities of the Marine Biological Laboratory (22°59’N, 113°97’E) at the Tsinghua Shenzhen International Graduate School. Groups of 10 corals were cultured in 120 L aquariums filled with artificial seawater (salinity = 33 ‰, pH = 8.2, T = 23-24 °C, light/dark = 12 h/12 h, Illumination with blue-dominated coral lamps in natural light, light intensity = 200 µmol m^-2^s^-1^). After allowing the corals to acclimatise to the laboratory environments for one month, we initiated our experiments.

### Symbiodiniaceae isolation

Symbiodiniaceae (*Symbiodinium* sp.) were fresh isolated from the *G. lobata*. First, the coral tentacles were clipped, and the microalgae were blown out repeatedly with a pipette gun. Second, the microalgae pellet was resuspended in seawater and centrifuged at 1000 g for 5 min at 20 °C. Third, the remaining host debris was removed using differential gradient centrifugation (900 g for 5 min → 700 g for 5 min → 650 g for 5 min at 20 °C)^85,86,87^. Finally, the microalgae pellet was centrifuged at 2000 g for 10 min at 20 °C and resuspended in F/2 medium, and then examined through an inverted microscope (Primo Vert, Germany) at 40× magnification. Isolated Symbiodiniaceae were enriched in F/2 medium for further experiments.

### Symbiodiniaceae carbon demand measurement

The carbon demand of Symbiodiniaceae was evaluted following the description by Wu et al.^40^ The seawater used in the experiment was decarbonised by the method of Yellowlees et al.^42^ and then added with corresponding concentrations of bicarbonate (NaHCO_3_). In detail, the pH was first adjusted to 4.5 with 1 M HCl to convert all DIC to CO_2_. Then, CO_2_ was removed by simultaneous aeration with N_2_ and agitation for 1 h. 1 mM Tris was then added, after which the pH was adjusted to 8.2 with 1 M of NaOH to ensure the pH stability. Different amounts of NaHCO_3_ were added to 40 mL-sealed headspace bottles containing 20 mL of artificial seawater medium as a carbon source, resulting in a final HCO_3_^-^ concentration of 2 mM, 4 mM, and 8 mM, respectively. An aliquot of 2 mL of Symbiodiniaceae culture (about 5.6×10^6^ cells mL^-1^) was added to each headspace bottle. Each dosage was triplicated independently. These experiments were conducted with continuous illumination (blue light, 200 µmol m^-2^s^-1^) at 23-24 °C^88^, and pH was maintained at 8-8.5^89^.

To ensure that DIC was the only variable and to exclude the effects of changes in N and P concentrations and light on microalgal growth, the incubation period was 36 h (Excluding other nutritional factors, i.e., N, P). To avoid CO_2_ diffusion to the air, the headspace bottles were sealed except during the sampling process, and a control group with only 22 mL of the corresponding concentration of NaHCO_3_ was added at each treatment to serve as a blank for the changes of DIC concentrations.

The biomass and DIC concentration in solution were measured every 12 h. Cell counts were performed with a hemocytometer under an inverted microscope (Primo Vert, Germany). Cellular dry weight was calculated by drying the microalgae in an oven at 55 °C for more than 24 h to keep cell weight constant. The standard curve between the cell counts and dry weight was established by preliminary experiments (Table. S2), and the dry weight of the biomass in this experiment was estimated based on cell counts. DIC concentrations were measured using a total organic carbon analyzer (Shimadzu, Japan), which detects dissolved CO_2_, HCO_3_^-^, and CO_3_^2-^ in solution. The average specific growth rate (g L^-1^d^-1^) over any two-time points was obtained by the following equation:

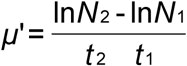

where *N*_2_ and *N*_1_ are the cell counts (g L^-1^) at time t_2_ and t_1_, respectively.

### Coral absorbance of organic and inorganic carbon under different light conditions

These experiments were carried out under two different light conditions: light (light/dark = 12 h/12 h, 200 µmol m^-2^s^-1^) and dark (continuous darkness), with or without organic carbon. Here, organic carbon is defined as decarbonised seawater with glucose (initial concentration of glucose 0.10 g L^-1^ based on the preliminary experiment) as the only carbon source, whereas inorganic carbon is defined as normal seawater with HCO_3_ ^-^ (initial concentration of HCO_3_ ^-^ 0.26 g L^-1^) as the main carbon source.

In total, four experimental treatments were set: DG (Continuous darkness and glucose as the only carbon source), DS (Continuous darkness and NaHCO_3_ as the only carbon source), LG (12 h/12 h light/dark and glucose as the only carbon source), and LS (12 h/12 h light/dark and NaHCO_3_ as the only carbon source). Each coral was cultured independently in 1.6 L of the respective medium. To exclude possible interference from stored sugars, all corals used had been starved in darkness for 3 days before experiments. Triplicates of each treatment were conducted During the experiment, coral visual changes were recorded every day using a camera at fixed points, and quantified by the mean values of tentacle extension length using a measuring ruler. The external DIC and glucose concentrations were detected simultaneously. Whole coral tentacles were ground and homogenised, and then used for measurements of carbon concentration and carbon fixation enzyme activities in coral tissues every other day. On day 8 of this experiment, NO_3_^-^ and PO_4_^3-^ remained in the medium.

### Coral bleaching induction by CO_2_ gas exposure

Two different species of scleractinian corals (*G*. *lobata* and *A. digitifera*) were selected for the coral bleaching experiment. The experiments were set up with three treatments: CO_2_ group (1% CO_2_ mixture gas was continuously aerated into seawater at 0.05 L min^-1^, with the microbubble plates generating microbubbles with an average size of 400-600 µm, pH = 7.0), HCl group (1 M HCl was added to seawater daily to adjust the same pH as the CO_2_ group to examine the possible effect of changes pH on the performance of coral, pH = 7.0) and control group (natural seawater, pH = 8.2), and other conditions were kept the same (salinity = 33‰, T = 23-24 °C, light intensity = 200 µmol m^-2^s^-1^, light/dark = 12 h/12 h). The same flow rate of air exposure did not cause the Symbiodiniaceae to leave, which can exclude the effects of bubbles’ physical disturbance on corals. Each coral was placed in a 1.6 L system and cultured independently. Each treatment was triplicated. During the experiment, the pH of seawater and CO_2_ solubility were measured daily. On the last day, free Symbiodiniaceae were collected from seawater, and symbiotic microalgae on corals were separated with water pick method. Meanwhile, the symbiotic microalgae coverage, photosynthetic activity (Fv/Fm), chlorophyII, carbon fixation enzymes, and anti-oxidative enzymes’ activities were detected. The dissolved CO_2_ were measured using a total organic carbon analyzer (Shimadzu, Japan), and its relationship with microbubble diameter was determined with the Planktonscope image system (Oasis Photobio Tech, China).

### Physiological parameters and enzymatic activities

DIC concentration in seawater was measured using a total organic carbon analyzer (Shimadzu, Japan) with the Chinese standard of HJ 501-2009, and glucose concentrations were measured by the dinitrosalicylic acid (DNS) method^90^, DIC and glucose consumption was calculated, respectively. A multi-excitation chlorophyll fluorometer was used to measure the photosynthetic activity (Fv/Fm) of microalgal cells. ChlorophyII was quantified using the methods of Ji et al.^91^ and Laura et al.^92^, respectively. Protein concentrations were determined with the bicinchoninic acid (BCA) method^93^ for homogenisation of all assay metrics. The activities of carbon fixation enzymes, i.e., CAs, PEPCase and PEPCKase, and anti-oxidative enzymes, i.e., SODase and CATase were measured by double antibody sandwich method using ELISA kits (Thermo Fisher Scientific, USA).

### Transcriptomics analysis

On day 8 of the coral culture experiments, all the corals were taken out and fast-frozen with liquid nitrogen for transcriptomic analysis. Total RNA was extracted using the Trizol-centrifugal column method, and then sent to the Shanghai Majorbio Bio-pharm Technology Co., Ltd. (Shanghai, China) for the construction of an RNA-seq transcriptome library. Each treatment was replicated three times. The data were analyzed using the Majorbio Cloud online platform (https://www.majorbio.com/). Raw reads were trimmed and quality controlled by fastp (Version 0.19.5).

To distinguish the sequences of coral and Symbiodiniaceae, the clean data from the samples were used to do *de novo* assembly with Trinity (Version 2.8.5) initially, and all the assembled sequencs were filtered by CD-Hit (Version 4.5.7) and TransRate (Version 1.0.3) to increase quality. Species annotation was performed through the NCBI-NR database to select coral species for subsequent analysis. Meanwhile, the clean reads were aligned to reference genome of Symbiodiniaceae (*Cladocopium goreaui*, http://sampgr.org.cn/index.php/download) with orientation mode using HISAT2 (Version 2.1.0), and the alignment rates ranged from 31.23% to 36.9%. The mapped reads were assembled by StringTie (Version 2.1.2).

The expression level of each transcript was calculated according to the transcripts per million reads (TPM) method. DEGs between four treatments were identified using DESeq2 (Version 1.24.0), and DEGs with |log_2_FC| ≥ 1 and P-value < 0.05 were considered to be significantly different expressed genes. Subsequently, DIAMOND (Version 0.9.24) was used to compare with the NCBI-NR database for species annotation, and with the KEGG database for functional annotation (E-value < 10^-5^, identity ≥ 50%, and comparison length ≥ 50%). Metabolic pathways were mapped using the KEGG Pathway as the primary reference. BLAST+ (Version 2.9.0) was used to compare with KEGG, Swissprot, and Pfam databases to obtain more coding genes and classification information of PEPC, PEPCK, and CA in the transcriptome.

### Statistical analysis and data availability

All experiments were conducted with a minimum of three biological replicates to ensure result reproducibility. The results are presented as mean ± standard deviation (SD). Statistical analysis was performed using SPSS (Version 26.0) and R (Version 4.4.0). Statistical significance was determined at thresholds of 0.05. All tested growth rates and activity indicators were analysed using two-way ANOVA, followed by Tukey’s HSD post-hoc analysis (*p* < 0.05). Genetic distances were calculated using MEGA11 based on the P-distance model, and phylogenetic trees were constructed using the Neighbor-Joining (NJ) method^94^. Heatmaps were drawn using ImageGP (https://www.bic.ac.cn/ImageGP/) and Z-Score normalisation was performed.

## Data availability

The data of this study are available within the article. Source data collected in this study are available from Zenodo online (https://doi.org/10.5281/zenodo.11276136). Raw RNAseq reads for differential gene expression analyses have been submitted to NCBI’s SRA database (http://www.ncbi.nlm.nih.gov) under BioProject PRJNA1115939. The KEGG database (https://www.kegg.jp/) was used for functional enrichment analyses.

## Code availability

All scripts for data visualisation are available from Zenodo online (https://doi.org/10.5281/zenodo.11276136).

## Acknowledgements

This work was supported by the Projects of Shenzhen Science and Technology Program (KCXFZ20211020163557022, KCXFZ20211020165547011), and the Cross Research and Innovation Funding of Tsinghua SIGS (JC2022004).

## Author contributions

BY.Z. and ZH.C. conceptualised and designed the experiments. BY.Z., L.L., MT.X., YQ.L. and WM.X. performed the experiments. BY.Z., S.T. and JM.Z. analysed and visualised the data. BY.Z. and S.T. interpreted the results and wrote the manuscript. HS.B., J.Z., M.C.B. and ZH.C. revised the manuscript.

## Supplementary Information

## Supplementary Information

**Fig. S1:**
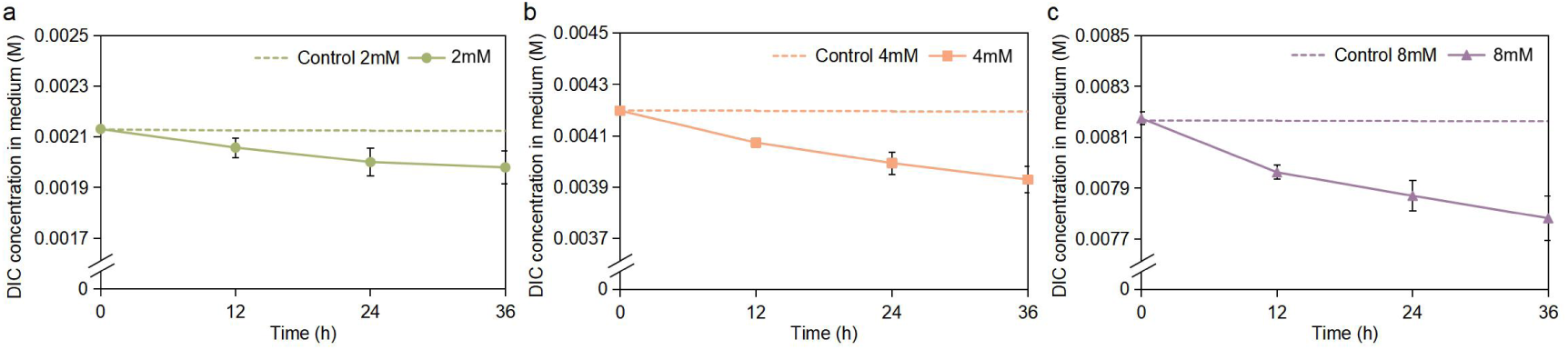
DIC consumption of monocultured Symbiodiniaceae at different carbon concentrations. **a-c**, DIC concentration in medium at 2 mM, 4 mM and 8 mM NaHCO_3_ group. The graphs show the mean ± SD (n = 3).

**Fig. S2:**
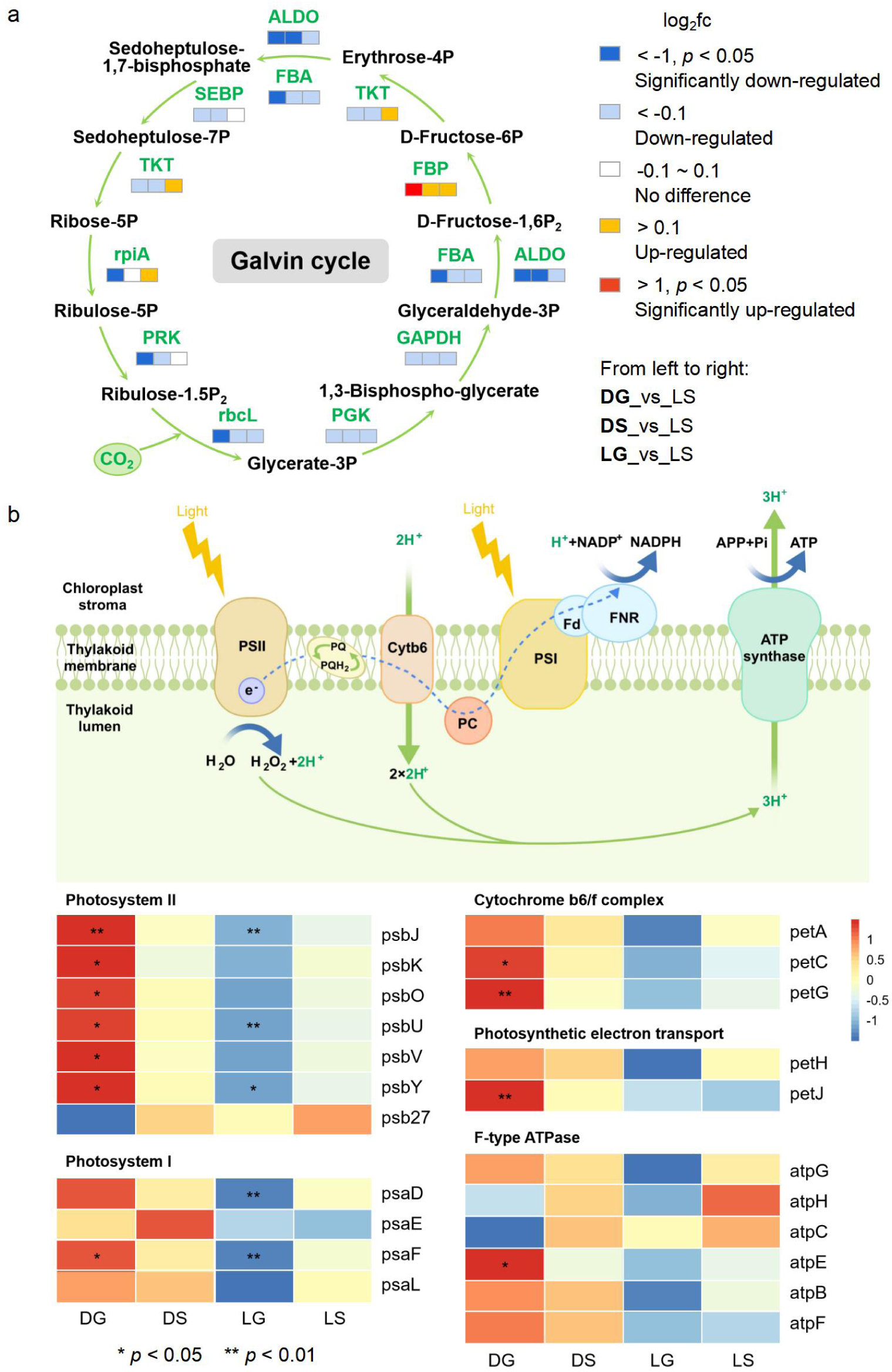
Comparison of the differential expression of photosynthesis genes in Symbiodiniaceae. a, Carbon fixation genes in Symbiodiniaceae. P_2_ bisphosphate. P phosphate. **b,** Photosynthesis photosystem genes in Symbiodiniaceae. The expression results of one sequence with the largest length of the transcriptome compared with the reference sequence of KEGG database were selected for each gene in the figure. The colors indicate the log_2_fc values of each treatment group compared to the control LS, from left to right once for DG, DS, and LG. Heatmap colors indicate relative expression levels, also the log_2_fc values of each treatment group compared with the control LS. P indicates the significance of the difference between each treatment group and LS.

**Fig. S3:**
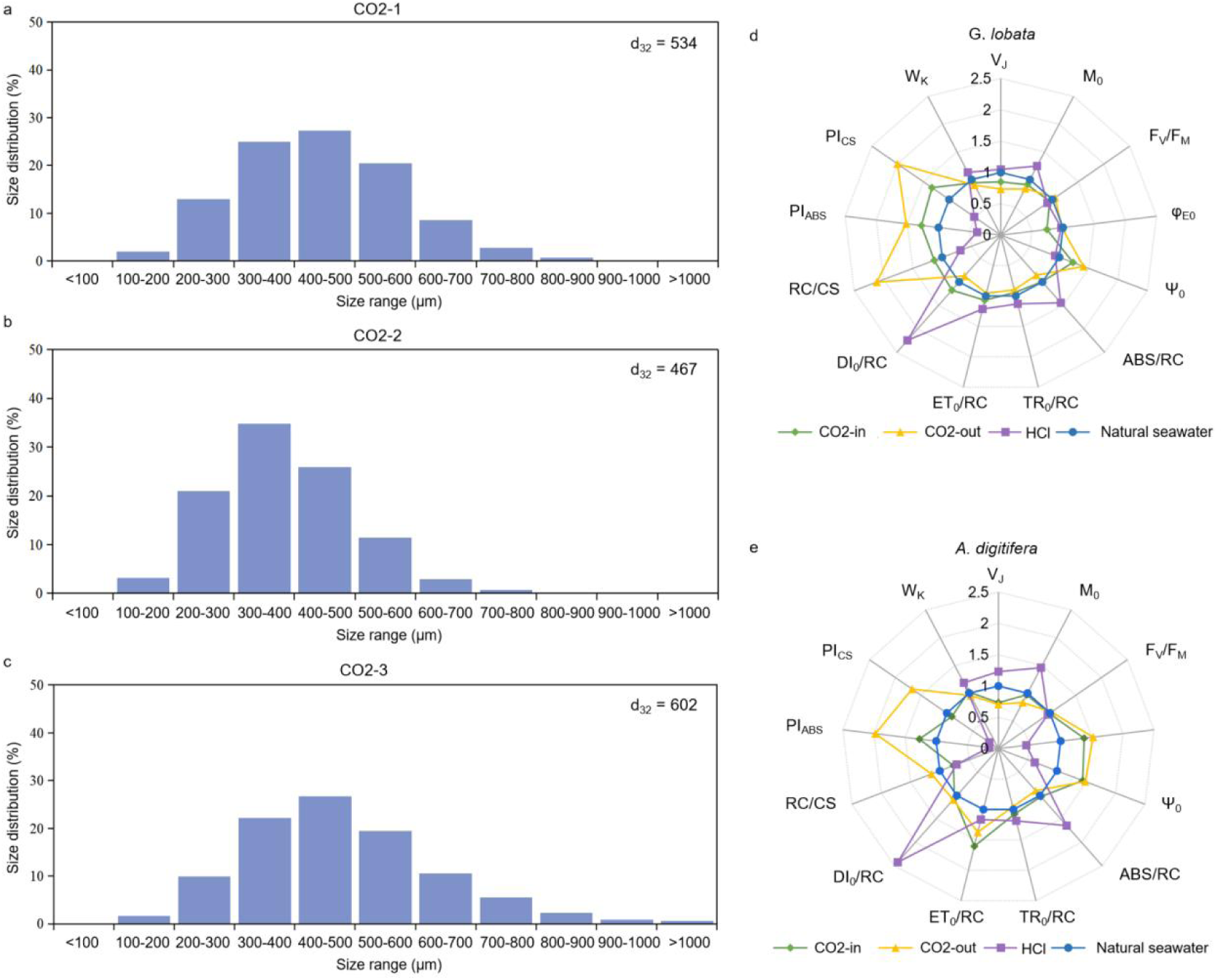
Size range of microbubbles in CO_2_ group and Chlorophyll fluorescence parameters of *G. lobata* and *A. digitifera*. **a-c**, Size range of microbubbles in CO_2_ group 1-3. d_32_ indicate the diameters of microbubbles. **d-e**, Chlorophyll fluorescence parameters of *G. lobata* and *A. digitifera*, respectively. The related parameters used in chlorophyll fluorescence quantification were shown in Table. S1.

**Fig. S4:**
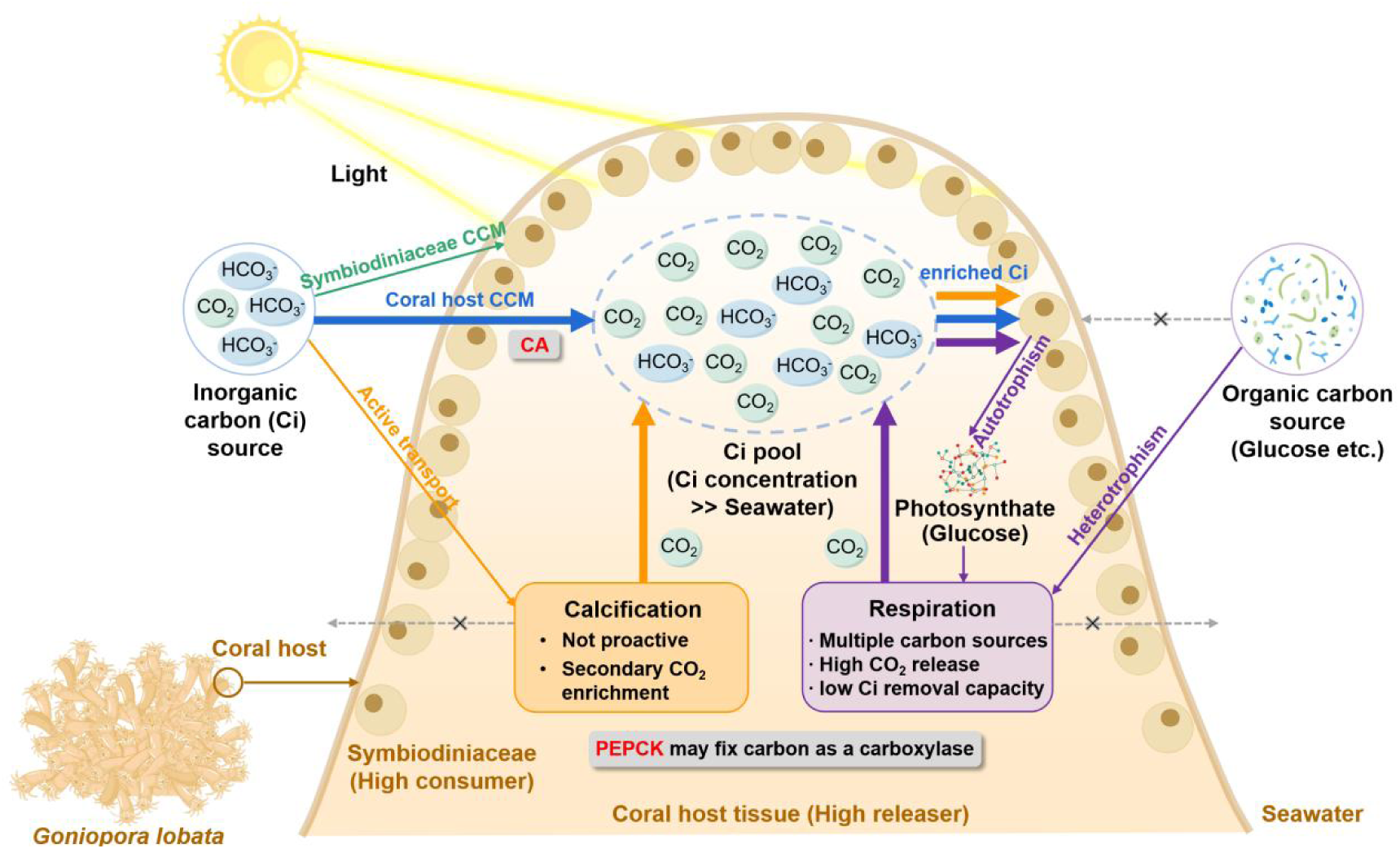
Schematic diagram of the carbon fixation pattern of coral host-Symbiodiniaceae symbiosis. The diagram indicates the internal tissue in one tentacle of the *G. lobata*. Arrow colors indicate different DIC pathways. Green indicates symbiotic algal CCM. Blue indicates host CCM. Yellow indicates host calcification. Purple indicates host respiration.

**Table. S1:**
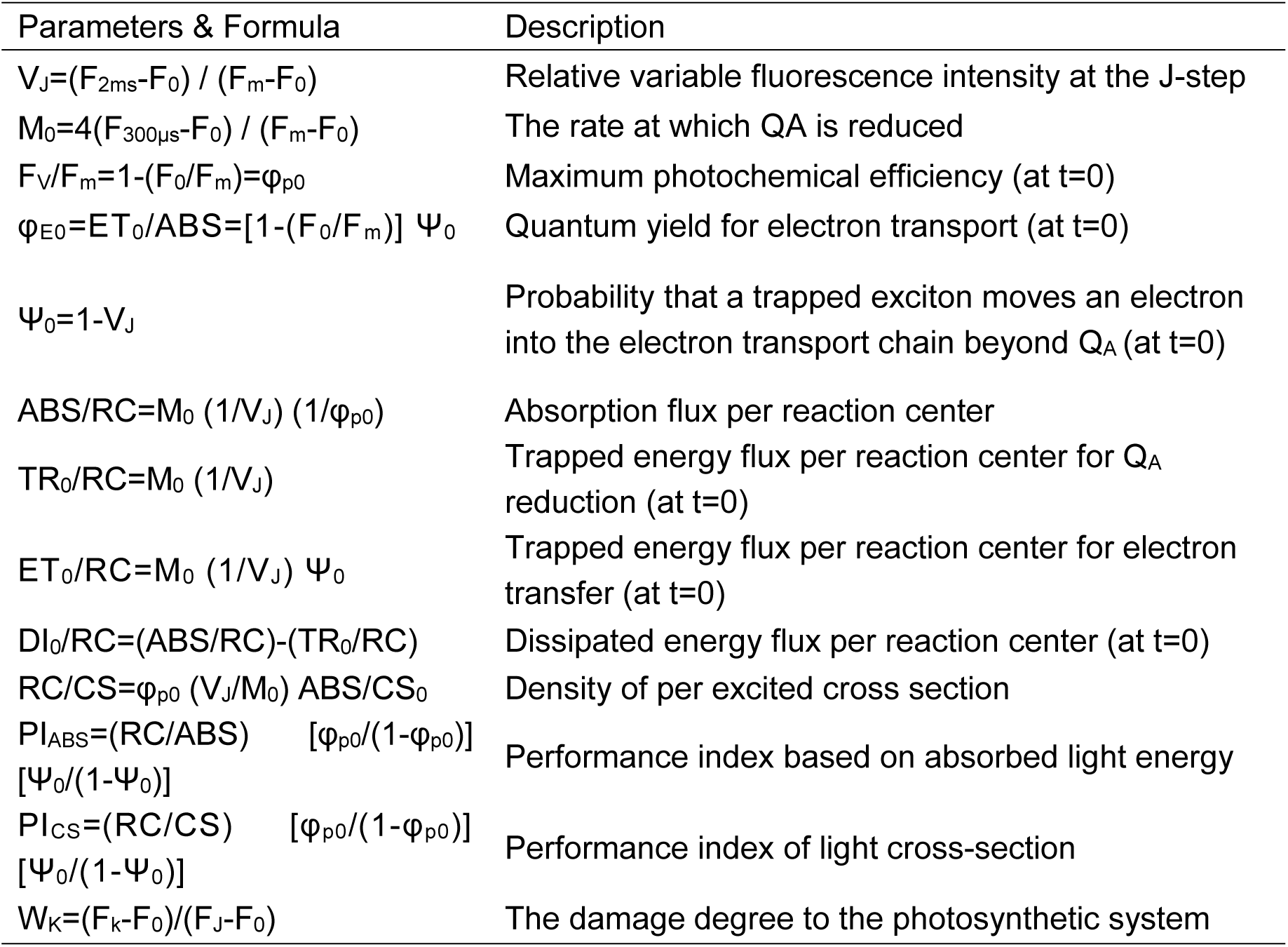
The related parameters, formula and related-descriptions used in chlorophyll fluorescence quantification.

**Table. S2:** Preliminary experiments to calculate the standard curve between the cell counts and dry weight of Symbiodiniaceae. Average = 4.05 ×10^-10^, SD = 1.12 ×10^-11^.

|  | Group 1 | Group 2 | Group 3 | Group 4 | Group 5 |
| --- | --- | --- | --- | --- | --- |
| Dry weight<br>(g mL<sup>-1</sup>) | 0.0031 | 0.0032 | 0.0029 | 0.0031 | 0.0032 |
| Cell counts<br>(cell mL<sup>-1</sup>) | 7.64 ×10<sup>6</sup> | 7.80 ×10<sup>6</sup> | 7.50 ×10<sup>6</sup> | 7.60 ×10<sup>6</sup> | 7.68 ×10<sup>6</sup> |
| Dry weight of<br>individual cells<br>(g cell<sup>-1</sup>) | 4.06 ×10<sup>-10</sup> | 4.10 ×10<sup>-10</sup> | 3.87 ×10<sup>-10</sup> | 4.08 ×10<sup>-10</sup> | 4.17 ×10<sup>-10</sup> |

